# Mechanical transapical coupling of endometrial epithelial cells during implantation

**DOI:** 10.1101/2023.12.12.571398

**Authors:** Jun Sakurai, Noriyuki Kinoshita, Tetsuhisa Otani, Hiroshi Koyama, Mikio Furuse, Toshihiko Fujimori

**Affiliations:** Division of Embryology, National Institute for Basic Biology, 5-1 Higashiyama, Myodaiji-cho, Okazaki 444-8787, Aichi, Japan; School of Life Science, The Graduate University for Advanced Studies, SOKENDAI, 5-1 Higashiyama, Myodaiji-cho, Okazaki 444-8787, Aichi, Japan; Division of Cell Structure, National Institute for Physiological Sciences, 5-1 Higashiyama, Myodaiji-cho, Okazaki 444-8787, Aichi, Japan

## Abstract

Epithelial cells lining the lumen of organs have an apical-basal polarity, and mechanical cell contacts are located in the lateral and basal regions. During early pregnancy in rodents, the luminal space of the uterus is closed, and opposing epithelial cells attach to each other at apical–apical surfaces. Here, we show that cells are mechanically coupled at the apical cell–cell contacts. The apical plasma membrane was intricately intertwined between opposing cells. Extracellular matrix and tight junction molecules were localized at apical contacts. Claudin inhibition resulted in impaired luminal closure, suggesting a functional requirement of claudins for the establishment of transapical coupling. The present results demonstrate a mechanical cell–cell interaction that occurs between apical–apical surfaces in addition to lateral and basal cell junctions.

**One Sentence Summaries:** Transapical coupling of epithelial cells mediated by extracellular matrix and tight junction molecules

## Introduction

Epithelial cells, which line the surfaces of the animal body and the lumen of organs, have a distinct apical–basal polarity. On the apical surface of epithelial cells, there are plasma membrane protrusions such as cilia and microvilli, which allow the transport of substances, and improve the efficiency of substance absorption and secretion (*1*). The integrin-dependent cell–substrate adhesion machinery or focal adhesions on the basal surface allow cells to attach to the basement membrane and extracellular matrix (ECM) (*2*) (*3*) (*4*). The cell–cell adhesion apparatus, including tight junctions, adherens junctions, and desmosomes, which mediate the mechanical interaction between neighboring cells, is located in the lateral domain (*5*) (*6*) (*7*). Transmembrane adhesion molecules that function in these adhesion complexes bind to cytoskeleton components forming a mechanically robust adhesion. Thus, the cell adhesion machinery identified to date is responsible for cell adhesion at sites other than the apical surfaces. However, there are situations in which the apical surfaces of epithelial cells are in close proximity in the luminal structure. For example, the apical surfaces of the retinal pigment epithelium and neural retina are in close apposition in the process of eye cup formation (*8*). The rodent uterus is also a luminal structure composed of a monolayer of endometrial epithelial cells or luminal epithelium (LE); the uterine lumen is closed and the apical surfaces of the LE are in close contact after the embryo reaches the uterus during implantation (*9*). Uterine luminal closure is critical for embryo implantation and pregnancy establishment, as embryonic development is suspended in the condition where the luminal space is kept open (*9*). The LE structures facing each other during uterine lumen closure are not well understood. This study examined the properties of the LE that change during implantation, and elucidated a new mode of mechanical cell–cell coupling at the apical surfaces or mechanical transapical coupling of interacting epithelial cells.

## Results

### A novel apical structure mediating mechanical coupling between the apical surfaces of two opposing epithelial cell layers

We examined the morphological changes of the LE of the endometrium during the implantation of mouse embryos. At 3.5 days after fertilization (E3.5), blastocysts are transported from the Fallopian tubes to the uterus and are evenly distributed along the uterine horns (*10*) in a fluid-filled luminal space. After 24 h, at E4.5, the apical surfaces of opposing LE cells are in close contact, and the uterine lumen is closed with no visible space (Fig. 1A). This occurs not only around the embryo, but throughout the uterus.

**Fig. 1.**
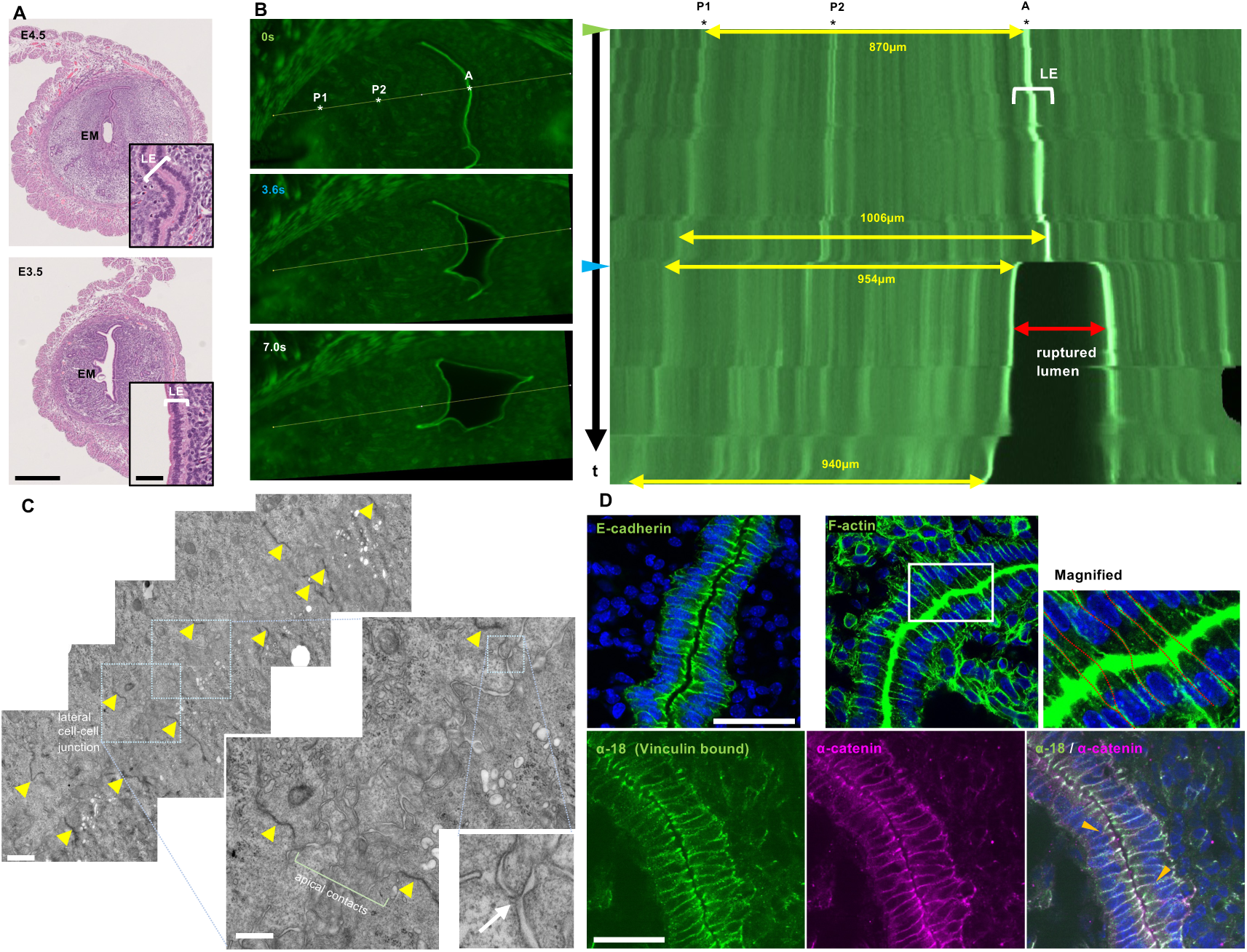
Mechanical coupling between the apical–apical contacts of epithelial cells during luminal closure in early mouse pregnancy. (**A**) Transverse section of the uterus at E4.5 and E3.5 stained with hematoxylin and eosin. Insets show magnified images. Scale bars: 500 µm (low magnification) and 50 µm (inset). EM; embryo, LE; luminal epithelium. (**B**) A 300 µm thick phalloidin-stained uterine section was expanded with forceps. The right panel shows a kymograph along the white line. The distance between the LE (indicated by “A”) and the bright point “P1” is shown by arrows. At 3.6 s (the blue arrowhead), the apical region was ruptured, and the luminal space became evident. (**C**) Electron microscopy of the apical–apical contact at E4.5. Left; low-magnification, middle; higher magnification images. Yellow arrowheads indicate lateral cell junctions with electron-dense signals. The lower right panel shows the triple bifurcation of the intercellular space (the white arrow.) Scale bars: 1 µm (left) and 500 µm (middle). (**D**) Sections showing E-cadherin, F-actin, α18-catenin, and α-catenin staining. The interconnection of lateral cell junctions between opposing epithelial cells is indicated by dashed red lines. Arrowheads indicate the V-shape branching of cell boundaries. Scale bars: 50 µm.

We hypothesized that the mechanical interaction between opposing cells at the apical surface ensures uterine luminal closure. To test this possibility, we applied an external force to separate opposing LE cells. We prepared a 300 µm thick transverse section of the pregnant uterus at E4.5 when the uterine lumen was closed, and separated the opposing epithelia with forceps. As shown in Fig. 1B and Movie S1, the stromal region (tissues including P1 and P2) was initially stretched when the force was applied, and the LE cells showed little change. However, after a certain amount of stretching at 3.6 s, a rapid change in the apical–apical contacts of LE cells was observed (as indicated by the blue arrowhead in the right panel of Fig. 1B). The space between the opposing epithelia widened, and this response spread along the LE cell sheets from the center as if the zipper or Velcro fastening had been opened. This suggests that the apical cell–cell contacts were ruptured under the applied forces. Meanwhile, the stretch of the stromal region was reduced, suggesting that the stromal region was stretched to buffer the applied force. The basal region of the LE attached to the surrounding stromal tissue was not broken, suggesting that the apical–apical contacts are weaker than those of other regions to allow mechanical separation. The abrupt rupture at the apical cell–cell contacts indicates that the contact site might contain a structure that is resistant to tensile forces. This observation supports our hypothesis that epithelial cells are mechanically coupled at the apical–apical contacts.

We then examined the structure at the apical contact site by transmission electron microscopy (Fig. 1C). Electron-dense adherens junctions at the lateral cell–cell adhesion sites in opposing cells were located at a distance of 1.2–1.6 μm. The apical plasma membrane of the opposing cells was intricately intertwined over an approximately 1 μm wide area. The distance between plasma membranes facing each other was <50 nm in the intertwined regions. Areas of triple bifurcation of intercellular spaces in which the plasma membranes were closer together were frequently observed (Fig. 1C, magnified image). These observations indicate that the structure at the apical regions between two opposing epithelia is different from the known apical structure. Microvilli containing actin bundles are present on the apical surface of endometrial epithelial cells when the uterine lumen is open (*11*). In the pregnant rodent, microvilli on the apical surface are lost in the process of “plasma membrane transformation” (*11*); however, our observations suggest that microvilli were transformed into the intertwined structure at the point of implantation. Objects resembling actin fiber are found in the cytoplasm of the intertwined structure, and the luminal closure is completed within less than 24 hours, which suggests that the newly found structure at apical cell–cell contact sites is established after the remodeling of the microvilli. We named this structure “transapical coupling”.

### Interconnection of lateral cell boundaries across the apical cell–cell contacts

Next, we determined the location of cell adhesion–related molecules in LE cells after the formation of the transapical coupling (Fig. 1D). Cell–cell adhesion sites containing E-cadherin were detected on the lateral sides, whereas E-cadherin was absent from the points of apical contact between cells, as observed in normal epithelial cells. F-actin was abundant at the apical contacts between epithelial cells as well as the lateral cell–cell adhesion regions. The line formed by lateral cell–cell adhesion regions was interconnected in some areas between opposing epithelial cells across the actin-enriched apical contacts (Fig. 1D, magnified image). Near the apical surface, some of the lateral cell boundaries diverged in a V-shape, suggesting that the number of cell boundaries between opposing cells are matched. Similar images of intercellular connection of lateral cell–cell adhesion sites were obtained with anti–α-catenin staining (Fig. 1D, lower panels). The monoclonal antibody α18 specifically recognizes α-catenin in a tension-transducing state (*12*). The α18 staining signal was enriched at the apical–lateral border in the region at which cell–cell adhesion sites face each other at E4.5, whereas at E3.5, it was uniformly present in the lateral side of the epithelium where the apical lumen was open (fig. S1). This suggests that the apical end of the lateral cell adhesion site was under greater tension after luminal closure, when opposing epithelial cell–cell adhesion sites were interconnected across the apical cell contacts.

### Mass spectrometry analysis detects ECM and tight junction components on the luminal surface of the uterus

E-cadherin and its cytoplasmic binding partner, α-catenin, were absent from the apical cell–cell contacts, suggesting that transapical coupling is mediated by mechanisms other than cadherin-mediated adhesion.

To identify the molecules involved in the formation of transapical coupling, we detected the proteins present at the apical surface of the endometrial lumen. We injected sulpho-NHS-biotin into the uterine lumen to biotinylate surface proteins. Visualization of the biotinylated areas with fluorescently labeled streptavidin showed that the apical surface of the epithelial cells of the uterine lumen was labeled (Fig. 2A). Proteins extracted from the uteri after biotinylation were trypsinized, and the biotin-labeled products were pulled down. Mass spectrometry analysis of the products showed that 98.7% of the peptides identified were biotinylated. GO analysis indicated enrichment of ECM-associated molecules (Fig. 2B). Samples from non-pregnant and pregnant E4.5 uteri were compared to identify proteins specifically present during uterine luminal closure. ECM proteins such as fibronectin, collagens, and laminins, as well as their receptor integrins were more abundant at E4.5 (Fig. 2C). In addition, differences in the tight junction proteins claudin-3, and claudin-10 were detected. Membrane proteins including receptors involved in intercellular signaling were also identified (data not shown). By contrast, mucin-1 and -4 were abundant in non-pregnant samples, suggesting that their expression decreases during pregnancy.

**Fig. 2.**
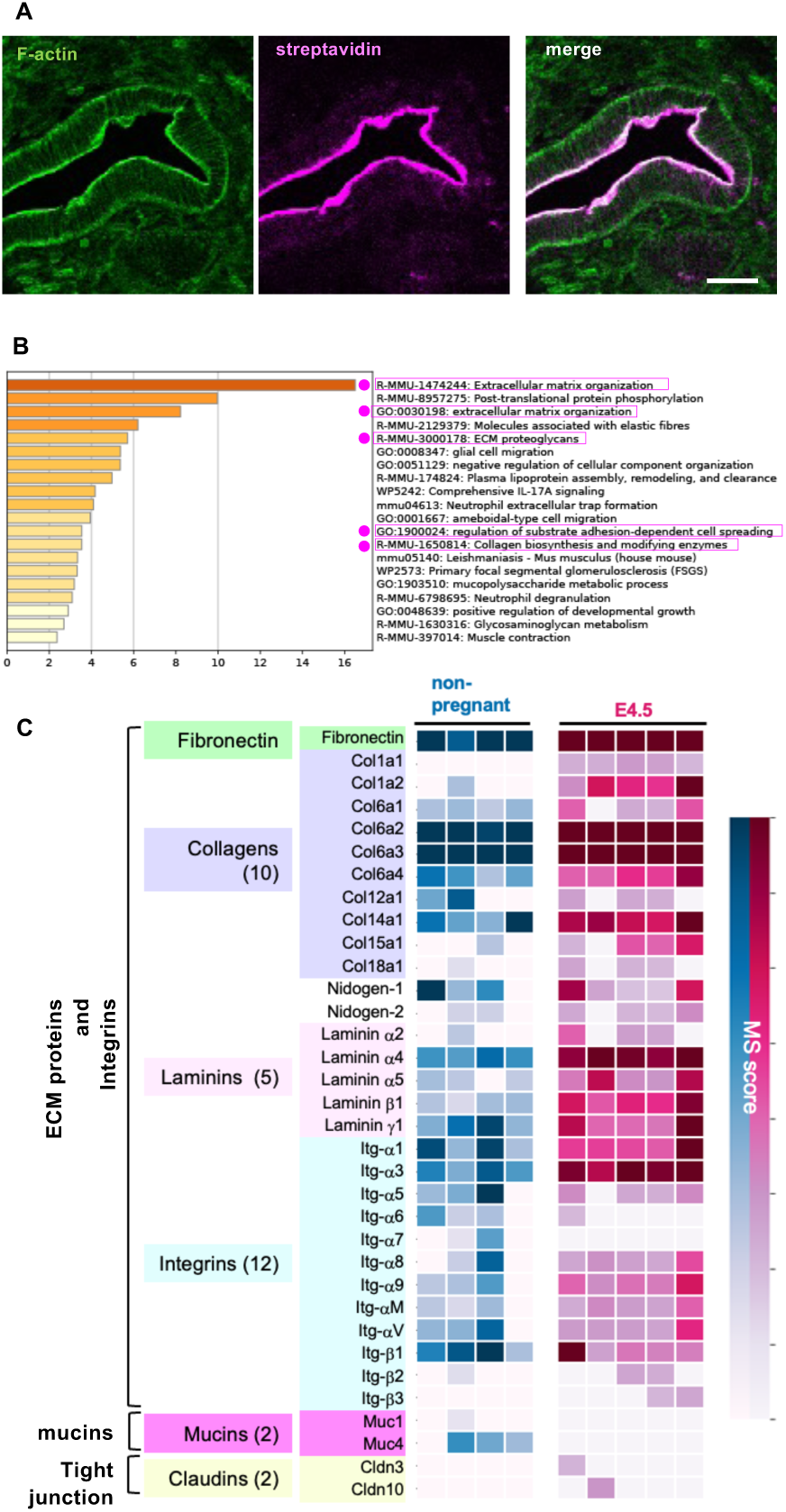
Identification of the apical surface molecules of LE proteins after biotinylation. (**A**) A biotinylation reagent, sulfo-NHS-biotin, was injected into the uterus dissected from a non-pregnant mouse. Biotinylation of the luminal surface was confirmed by staining uterine sections with fluorescence-labeled phalloidin and streptavidin. Scale bars: 20 µm. (**B**) GO analysis of biotinylated proteins commonly identified in nine independent samples by mass spectrometry (52 proteins). The analysis was performed in Metascape (https://metascape.org/). GO terms related to the extracellular matrix (ECM) were enriched, as indicated by magenta circles and boxes. (**C**) Normalized mass spectrometry scores of ECM-related proteins, mucins, and tight junction proteins are indicated in the heatmap. Each column represents independently prepared samples for mass spectrometry: four samples from two non-pregnant mice are colored in blue, and five samples from three E4.5 mice are colored in red.

### Integrin and ECM components are present on the apical surface

Among the proteins detected by mass spectrometry (Fig. 2C), we initially focused on the ECM components and integrins. To determine whether these proteins were located on the apical surface of the LE, we performed immunofluorescence detection of integrin-α5, −α1, and −αV, fibronectin, collagen (Col1A2), and laminin, and the localization of these proteins was compared between E4.5, E3.5, and non-pregnant stages (Fig. 3A). Integrin-α5 signals were detected at apical–apical contacts without any gaps at E4.5, when the lumen was closed, in addition to weak signals detected on the basal side of the epithelium. Integrin-α5 was not detected in LE cells in non-pregnant samples, and it was already present on the apical region at E3.5 before luminal closure. These observations suggest that the characteristics of LE cells changed under the hormonal control of the pregnancy. As shown in Fig. 3A, integrins-α1 and −αV, fibronectin, and Col1A2 also accumulated on the apical contacts between the opposing LE at E4.5. Fibronectin is a ligand for integrin-α5β1 and integrin-αVβ1, and collagen is a ligand for integrin-α1β1 (*13*), which is consistent with our observation that these proteins were present at the transapical coupling site. At E3.5, these proteins were localized at apical and lateral cell junctions or cytoplasmic vesicles, and were barely detected in LE cells under non-pregnant conditions. By contrast, laminin was detected mainly on the basal side regardless of the maternal conditions, suggesting that the LE maintained the basal membrane structure, and the apicobasal polarity was not disturbed, despite the appearance of ECM components on the apical side. The ECM components and integrin subunits accumulated before luminal closure during early pregnancy and could be involved in the initiation and maintenance of transapical coupling. We also examined the location of the kinases FAK (focal adhesion kinase) and ILK (integrin-linked kinase), which are associated with intracellular integrin complexes (*13*). These kinases also localized to the apical contacts at E4.5 (Fig. 3B and C). The localization of these kinases to the apical side together with some integrins and the ECM proteins indicates that the integrin complexes may function in transapical coupling.

**Fig. 3.**
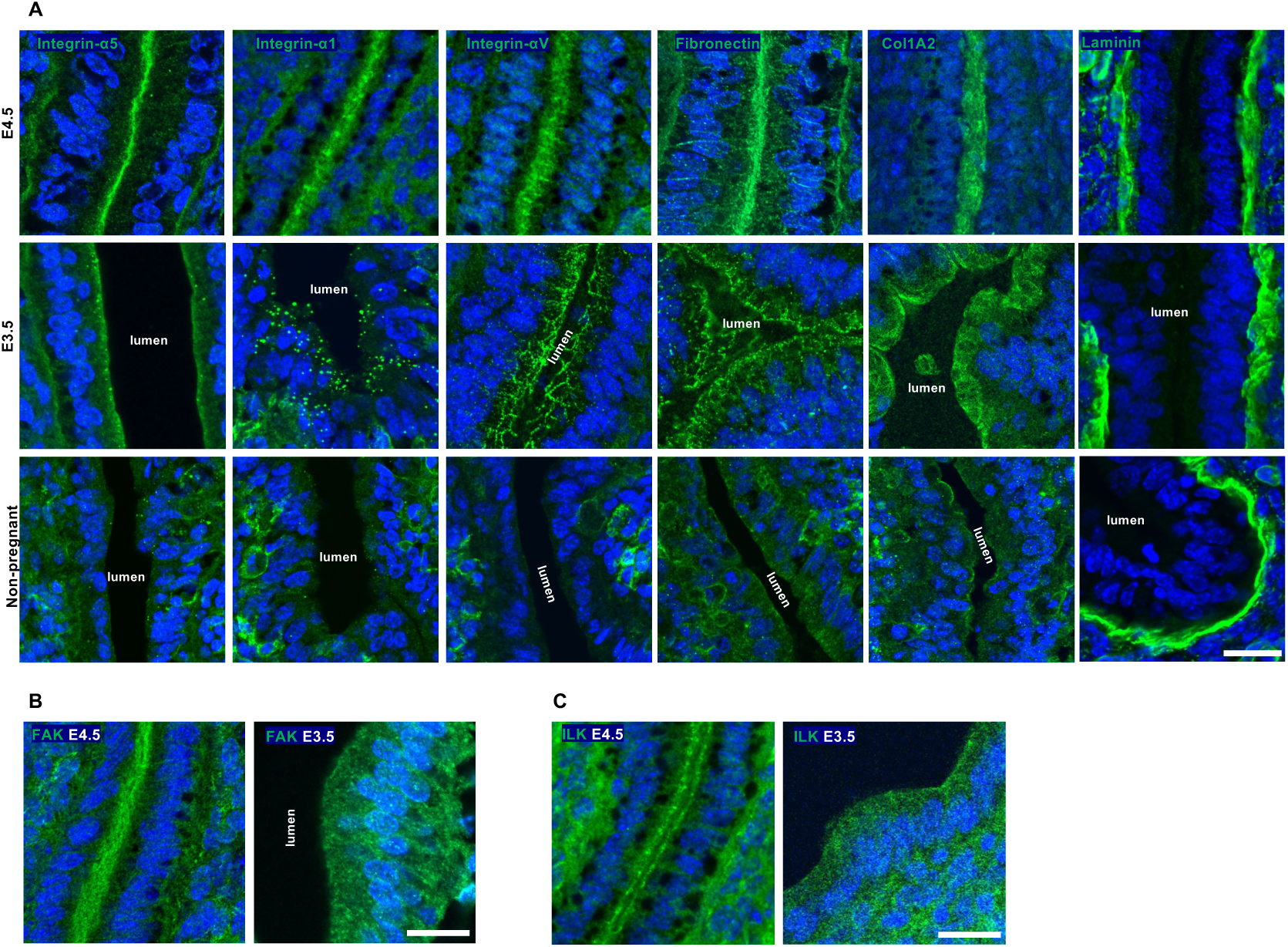
Accumulation of integrins and ECM proteins on the apical surface of the LE. (**A**) Immunostaining of transverse sections from E4.5, E3.5, and non-pregnant uteri using antibodies against integrin-α5, −α1, and −αV, fibronectin, collagen type 1A2 (Col1A2), and laminin during pregnancy. (**B**) and (**C**) Localization of focal adhesion kinase (FAK) and integrin-linked kinase (ILK) at E4.5 and E3.5. Scale bars: 20 µm.

### Tight junction protein, claudin-3, appears on the apical contacts

Because claudin-3 and claudin-10 were detected at E4.5 in the mass spectrometry analysis (Fig. 2C), we next performed immunofluorescence of tight junction proteins. Normally, tight junction proteins are found at lateral cell–cell junctions and concentrated especially at the apical–lateral boundary of epithelial cells (*7*). As shown in Fig. 4A, claudin-3 was detected at the apical contacts at E4.5 after luminal closure. Claudin-3 was distributed along the lines in the apical contact regions, as well as at the lateral cell–cell junction. The unique distribution of claudin-3 at the apical contacts suggests that this transmembrane adhesion molecule is involved in the transapical coupling mechanism. On the other hand, cytoplasmic tight junction protein ZO-1 localized to the apical–lateral boundary of epithelial cells. Analysis of other tight junction proteins showed that JAM-A (Junctional Adhesion Molecule-A) (*7*), and the tricellular junctional transmembrane proteins tricellulin (MarvelD2) and angulin-1 [lipolysis-stimulated lipoprotein receptor (LSR)] (*14*) localized to the apical–lateral boundary of epithelial cells while extended to the apical region compared to ZO-1 signals (Fig. 4B and fig. S2). Tricellulin and angulin-1 were located at areas of triple bifurcation extending from the lateral cell–cell junction. And signals for tricellulin and angulin-1 were partially overlapped with the signals for ZO-1, which may correspond to the branching points of lines of ZO-1 signals (marked with arrowheads in the Fig. 4B). These findings suggest that the tight junction proteins at the apical–lateral border are expanded for the formation of transapical coupling.

**Fig. 4.**
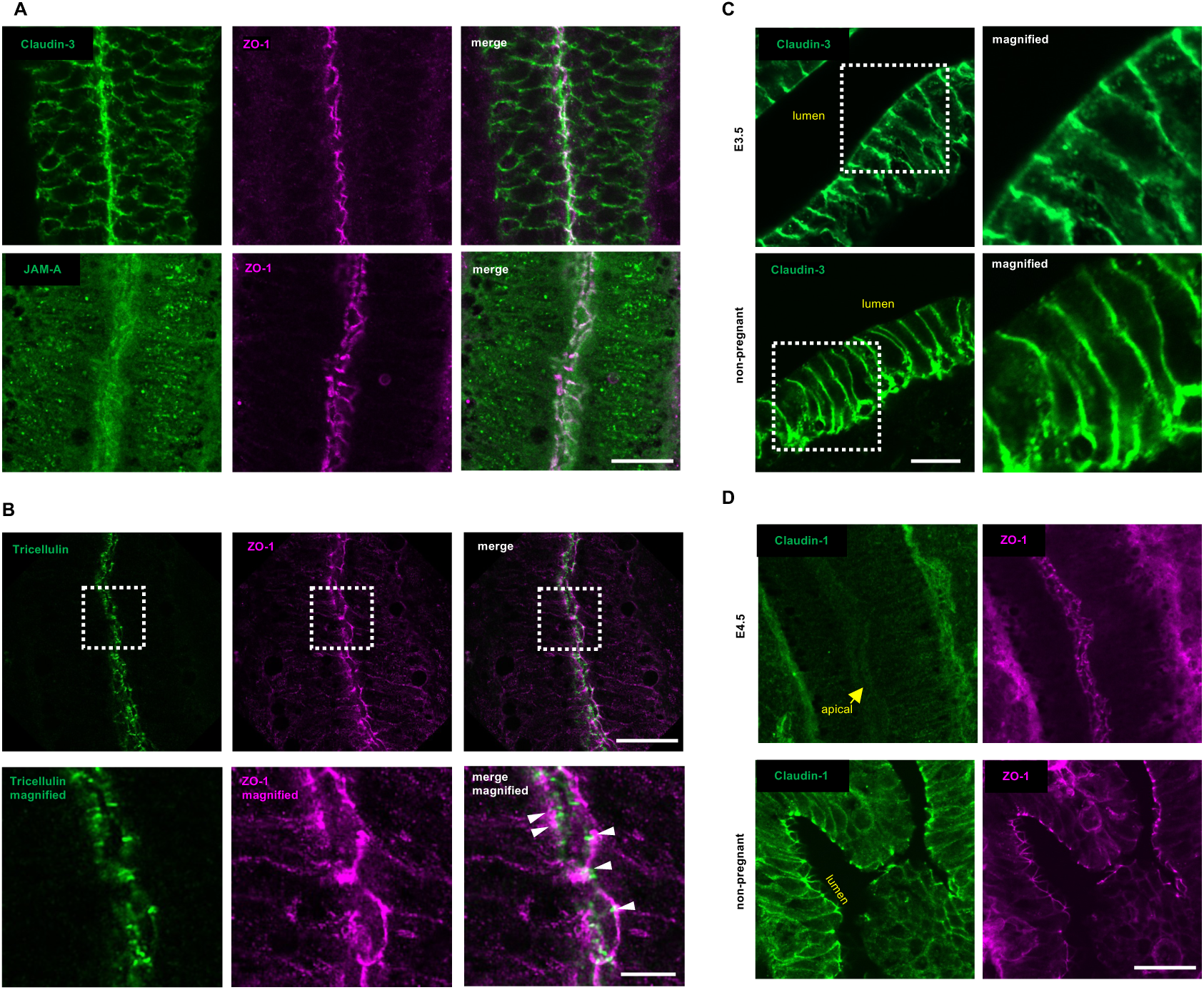
Localization of tight junction components in the apical contacts. (**A**) Localization of claudin-3, ZO-1, and JAM-A at the apical contacts of the closed LE at E4.5. Scale bars: 20 µm. (**B**) Localization of tricellulin in the apical contacts. Scale bars: 20 µm.The bottom panels show magnified images Scale bars: 5 µm. White arrowheads indicate the tricellulin signals that overlap with the branch points of the ZO-1 lines. (**C**) Changes in claudin-3 localization during pregnancy. Immunostaining of E3.5 and non-pregnant uteri. The left panels show magnified images. Scale bars: 50 µm. (**D**) Localization of claudin-1 during pregnancy. E4.5 and non-pregnant uteri were immunostained with claudin-1 and ZO-1 antibodies. The apical contact is indicated by a yellow arrow. Scale bars: 20 µm.

Next, we examined the timing of claudin-3 appearance on the apical surface. As shown in Fig. 4C, claudin-3 was enriched at the apical–lateral boundary, and weak signals were detected at the apical plasma membrane at E3.5 in addition to the lateral region. In the non-pregnant uterus, claudin-3 localization was limited to the lateral cell– cell junction with no accumulation around the apical region. These observations indicate that the localization of claudin-3 is dynamically regulated during gestation. Immunofluorescence detection of claudin-1 showed that it was enriched at the apical– lateral boundaries in addition to the lateral cell–cell junctions in the non-pregnant uterus; however, its expression was weak at E4.5, whereas that of ZO-1, a tight junction cytoplasmic protein, was maintained throughout all stages (Fig. 4D). These observations suggest that although the tight junction is maintained during pregnancy, the proteins mediating junction formation switch from claudin-1 to 3, and the localization of claudin-3 is expanded to the apical region when transapical coupling is established.

### Inhibition of claudin function impairs transapical coupling formation

To test whether the claudins at the apical contacts mediate the transapical coupling between LE layers, we inhibited claudin function using a C-terminal polypeptide of *Clostridium perfringens* enterotoxin (C-CPE) that binds to the extracellular loop of claudin-3 (*15*) and inhibits tight junction strand formation (*16*). C-CPE fused with glutathione S-transferase (GST) (Fig. 5A) was administered to the pregnant female at the E3.5 stage, and LE closure was examined at E4.5. Injection of control GST did not have detectable effects. However, injection of C-CPE inhibited LE closure in some regions (Fig. 5B), and luminal spaces were observed between opposing LE cells (Fig. 5C). In areas in which the apical regions of the LE were separated, the apical localization of claudin3 was decreased, and the ZO-1 signals were absent (Fig. 5B). These observations suggest that the functions of tight junction proteins are required for transapical coupling formation.

**Fig. 5.**
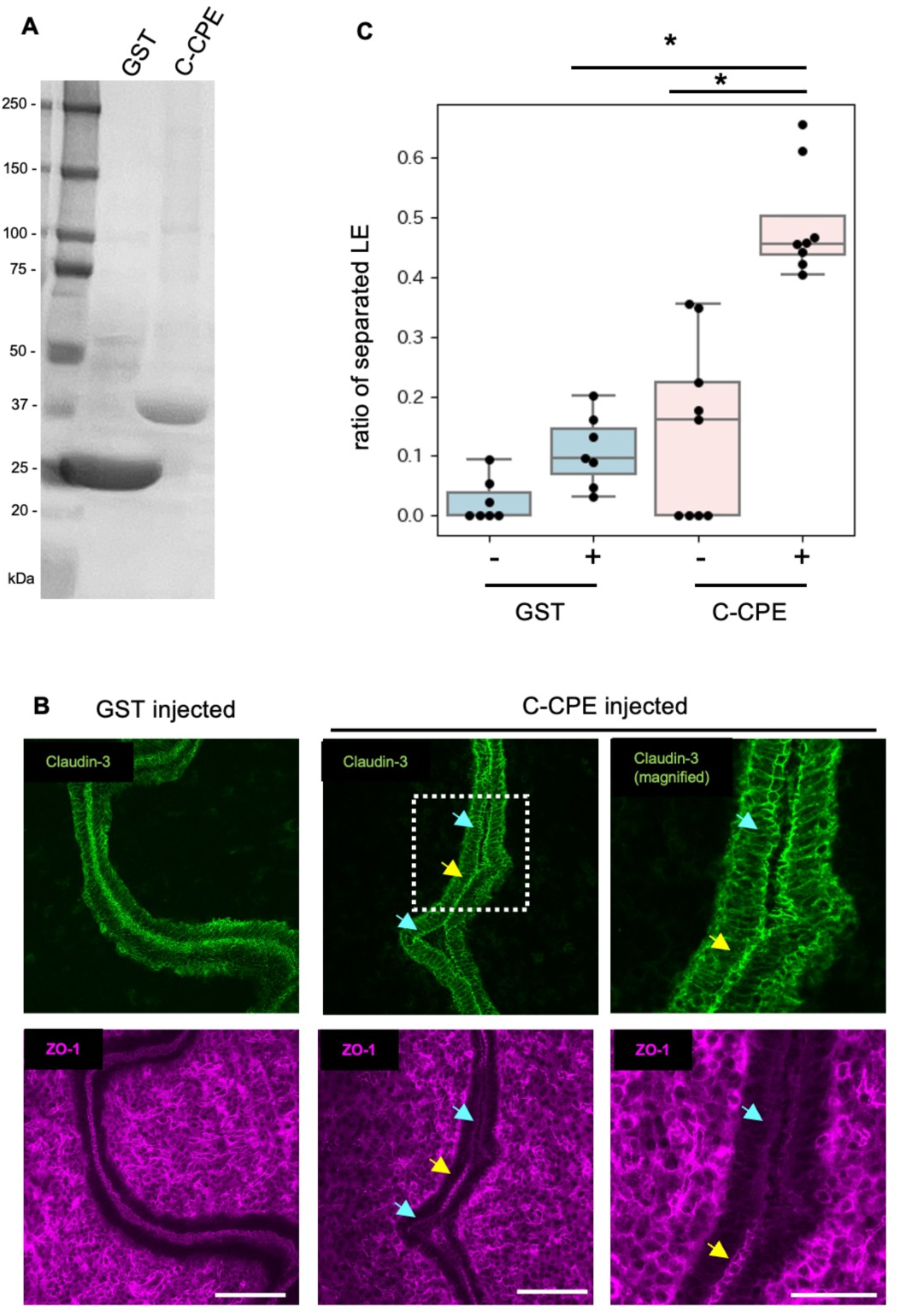
Claudin is required for transapical coupling formation. (**A**) Purified GST or GST-tagged C-CPE was separated by SDS-PAGE (silver-staining). (**B**) Effects of C-CPE, which inhibits the function of claudins, on luminal closure. Purified GST-tagged C-CPE or GST (shown in A) was injected into the lumen of the E3.75 uterus, and LE regions were examined at E4.5. Images of specimens immunostained using claudin-3 and ZO-1 antibodies. Yellow arrows, apical contact regions; blue arrows, separated regions. Scale bars: 100 µm in the left and middle panels and 50 µm in the magnified right panels. (**C**) The ratio of separated LE length to the whole LE length was plotted. *: p < 10^−4^ in samples without (−) or with (+) injection of purified proteins. Each dot represents a section.

## Discussion

### Mechanical transapical coupling of epithelial cells

The above results suggest that the apical surfaces of the mouse endometrial LE physically interact and are mechanically coupled with each other during luminal closure at implantation through a novel mechanism that we termed “transapical coupling”. The mechanical separation experiment suggested that the mechanical coupling was weaker on the apical contacts than on the basal and lateral regions, in which classical cell adhesion is observed (Fig. 1B). Mechanical transapical coupling allows uterine luminal closure by active function of the luminal epithelium rather than passive compression by surrounding tissues such as stroma and smooth muscle. The molecular characteristics of the apical region of epithelial cells changed during the establishment of transapical coupling (Figs. 2, 3, and 4). The mucous barrier proteins mucin-1 and -4 disappeared from the apical region (fig. S3), and extracellular matrix proteins and their receptor integrins were enriched in the apical region during early pregnancy. Tight junction protein, claudin-3, was also detected in the apical-apical contacts, regulating mechanical coupling formation (Fig. 5). The structure of the transapical coupling was different from that of the normal tight junction for the following reasons. Plasma membrane kissing points, where adjacent plasma membranes appear to be in close contact, which is one of the properties of the tight junction, are not evident in the transapical coupling. Cytoplasmic protein, ZO-1 is not always colocalized with claudin-3 at apical contacts. And the ECM proteins are also found. This suggests that the transapical coupling is a novel structure composed of ECM and tight junction components. These results indicate that epithelial cell–cell junctions are not limited to the lateral and basal regions, but are also evident in the apical region in certain cell types.

Physical interactions between the apical surfaces of epithelial cells are also observed during neural tube closure in vertebrates and chordates. In this process, the apical tips of the left and right neural fold cells eventually fuse to form the neural tube (*17*) (*18*). Previous studies revealed the presence of cellular protrusions from the apical region of the closing neural tube (*19*) induced by small GTPases. We examined the mouse closing neural tube at E8.5, but did not find structures similar to those detected in the uterine epithelium (data not shown). Close positioning of the apical surface of the epithelium also occurs during the formation of the vertebrate retina, when the lumen of the eye cup is closed, resulting in direct contact between the neural retina and retinal pigment epithelial (RPE) cells. In this case, the microvilli of RPE cells interact with photoreceptor outer segments of the neural retina (*20*). Another example is the fusion of three prominences of the facial epidermis during the formation of the lips, mouth, and palate, in which a disturbed process results in a cleft of the lip or the palate (*21*) (*22*) (*23*). However, there are currently no reports of similar structures observed in these tissues. A similar F-actin–rich structure with intertwined plasma membrane was reported between the apical surfaces of stalk cells in the ovariuterine epithelium of the scorpion *Euscorpous italicus*, although the molecular mechanisms underlying the formation of this structure are not known (*24*).

### What is the significance of transapical coupling during the establishment of embryo implantation?

In rodents, embryos evenly spaced along the longitudinal axis of the uterus settle in the implantation chamber to begin implantation (*10*) by interactions between the embryonic trophectoderm (TE) and the endometrial epithelium. The closure of the uterine luminal space after the formation of transapical coupling may restrict the movement of the embryo within the uterus to initiate implantation. Indeed, when embryonic diapause or delayed implantation is induced in mice, the luminal space is open even when the embryos hatch from the zona pellucida. This suggests that luminal closure after transapical coupling is a critical step in embryo implantation and establishment of the pregnancy.

## Acknowledgements

This work was supported by the Trans-Omics Facility and Optics and Imaging Facility, NIBB Trans-Scale Biology Center. This work was supported by the NIBB Collaborative Research Program 22NIBB410 and 23NIBB408. We thank Yumiko Makino and Tomoko Mori for the mass spectrometry operation. We also thank all the members of the Division of Embryology, NIBB for their advice and discussion.

## Author Contributions

Conceptualization: JS, NK, TO, MF, TF

Methodology: JS, NK, TO, TF

Data analysis: JS, NK, HK, TF

Funding acquisition: JS, NK, TO, TF

Project administration: NK, TO, MF, TF

Supervision: TF

Writing: JS, NK, TO, HK, MF, TF

## Data and materials availability

All data are available in the manuscript or the supplementary materials.

## Supplementary Materials

Materials and Methods

figs. S1 to S4

References (*25–33*)

Movie S1

