## Supplementary Materials for "Mechanical transapical coupling of endometrial epithelial cells during implantation"

**The PDF file includes:**

Materials and Methods
Figs. S1 to S3
References

**Other Supplementary Materials for this manuscript include the following:**

Movie S1

**Materials and Methods**

References 25-31 are only cited in the SM.

**Animals**

Mice (Slc; ICR, Japan SLC) were used for analysis. Female mice (10–28 weeks) were selected randomly and were crossed with males to induce pregnancy. Noon on the day of vaginal plug detection was designated as E0.5 (embryonic day 0.5). Animals were maintained in a light- and temperature-controlled room with a 12 h:12 h light:dark cycle at 23 ± 2°C. Animal care and experiments were performed according to the guidelines of Animal Experimentation of the National Institutes for Natural Sciences. All animal experiments were approved by Animal Research Committee of National Institutes for Natural Sciences (23A030).

**Preparation of hematoxylin and eosin–stained serial sections**

Whole mouse uteri were dissected from mice and fixed in 4% paraformaldehyde (PFA) for 2 days at 4°C. After the fixation, the tissues were washed three times with PBS, dehydrated in 70%, 90%, and 100% EtOH, and then immersed in chloroform, 100% paraffin wax (Paraplast plus, Tyco) at 62°C, and embedded in paraffin. The tissue blocks were sectioned at 5 μm using a microtome with flow function (HM340E, Microm), and the sections were placed on uncoated slides (S2444, Matsunami). Hematoxylin and eosin staining (Hematoxylin, Muto; Eosin, Muto) was performed using an automatic staining machine (Autostainer XL, Leica), and samples were mounted in MGK-S (Matsunami) with coverslips (Matsunami). All serial sections were scanned with a slide scanner (SCN400, Leica) and observed with a viewer (Aperio ImageScope, Leica).

**Examination of the mechanical interaction between opposing LE cells**

Pregnant ICR uteri at E4.5 were dissected and fixed in 4% PFA overnight at 4°C. The whole uteri were then washed with PBS, and transverse sections of 300 μm thickness were prepared using a vibratome (VT1200, Leica). Uterine sections were stained with Alexa Fluor 488 phalloidin (Invitrogen) to visualize the cell–cell borders. Under a stereomicroscope (SMZ25, Nikon) equipped with a camera (DS-Fi3, Nikon), force was applied with forceps to separate the opposing LE. Changes in the uterine tissues were recorded using NIS-Elements software (Nikon) at 30 ms intervals. The recorded time-lapse data were registered to reduce temporal translation of the images using ImageJ/Fiji software (a plugin "Linear Stack Alignment with SIFT" was used) and used to generate the kymograph with ImageJ/Fiji (a function "analyze > Multi kymograph" was used).

**Transmission electron microscopy**

Uteri were dissected from ICR E4.5 mice, separated into decidua units, and fixed in 4% PFA in 0.1 M phosphate buffer, pH 7.4 for 2 h at room temperature (RT), followed by 2.5% glutaraldehyde/2.5% PFA in 0.1 M phosphate buffer for 2 h at RT, and then transferred to 4°C. After fixation, the uteri were washed with 0.1 M phosphate buffer and sectioned at 150 µm thickness using a vibratome (VT1200, Leica) in phosphate buffer to expose the LE around the implantation site. All reagents used for sampling were filtered through a 0.22 µm filter. Uterine sections were postfixed with 1% OsO4 in 0.1 M cacodylate buffer, pH 7.4, for 1 h on ice. After washing with water, the specimens were stained with 0.5% uranyl acetate for 2 h at RT. After washing with water, sections were dehydrated in a graded series of ethanol. The sections were then immersed in propylene oxide for 1–2 min, transferred to a 1:1 mixture of propylene oxide and Quetol 812 resin, and incubated overnight. Sections were embedded in Epon 812 resin. Semi-thin sections of 500 nm were cut and stained with toluidine blue to examine the sample preparations. After further trimming of the section, ultrathin sections (50–80 nm) were cut and mounted on 200-mesh formvar-coated copper grids. The specimens were observed with a

JEM1011 or JEM1010 transmission EM (JEOL) at an accelerating voltage of 80 kV. Images were captured with a MegaViewG2 or Veleta CCD camera using iTEM software (all from Olympus Soft Imaging Solutions). Image processing (brightness and contrast adjustment) was performed using ImageJ software.

**Immunohistochemistry**

To visualize the implantation site, 1% Evans blue dye was injected from the tail vein. Uteri were dissected from mice and fresh frozen in OCT compound (Sakura-Finetek) or fixed in 4% PFA (4°C overnight), methanol (−20°C, 60 min), and acetone (−20°C, 5 min). After fixation, the uteri were washed three times with PBS and placed in 15% sucrose. After 6 h the uteri were incubated in 30% sucrose overnight and embedded into the OCT compound. For immunohistochemistry, 12–20 μm thick frozen sections were washed with PBS containing 0.1% Triton X-100 (PBST) and then incubated with Blocking One (Nacalai Tesque) at RT for 1 h to block non-specific binding of antibodies. Then, sections were incubated with primary antibodies in Blocking One at 4°C overnight. After washing with PBST, the sections were incubated with secondary antibodies at RT for 2 h or at 4°C overnight. After washing, samples were mounted in Fluoromount-G (Southern Biotech).

Fluorescence images were captured using a Nikon A1 confocal microscope or a Nikon Ti2 equipped with a spinning disc confocal scanner unit (CSU-W1, Yokogawa) and an S-CMOS camera (ORCA-Fusion BT, Hamamatsu Photonics).

Anti-ECCD2 (diluted 1:1) (*25*) antibody-producing hybridoma was a gift from Masatoshi Takeichi, RIKEN BDR, Japan. Anti–ZO-1 (diluted 1:1) (*26*), anti-tricellulin (diluted 1:1) (*27*) , and anti–angulin-1 (diluted 1:1) antibodies (*28*) were stocked at Mikio Furuse's lab. Anti–α18 catenin (diluted 1:1) antibody (*29*) was a gift from Akira Nagafuchi, Nara Medical University,

Japan. The antibodies against the following proteins were purchased: α-catenin (1:1000, C2081, Sigma-Aldrich, RRID: AB_476830), integrin-α5 (1:100, BS-0567R-TR, Bioss Antibodies), integrin-α1 (1:100, PAB313MU01, CLOUD-CLONE COUP.), integrin-αV (1:100, BS-2250R-TR, Bioss Antibodies), fibronectin (1:100, GTX102996, GeneTex,), Col1A2 (1:100, GTX102996, GeneTex), laminin (1:100, ab11575-250, Abcam,), claudin-3 (1:100, 341700, Thermo Fisher Scientific), claudin-1 (1:100, 519000, Thermo Fisher Scientific, RRID: AB_2533916), JAM-A (1:100, 36-1700, Thermo Fisher Scientific, RRID: AB_2533241), ZO-1 (1:100, 33-9100, Thermo Fisher Scientific, RRID: AB_2533147), FAK (1:100, HPA001842, Atlas Antibodies, RRID: AB_1855891), ILK (1:100, HPA048437, Atlas Antibodies, RRID: AB_2680391), Muc1 (1:500, 102352-T10, SinoBiological),. Alexa Fluor–conjugated secondary antibodies (1:500; Thermo Fisher Scientific), and Hoechst 33258 (1:100,000; Molecular Probes, OR, USA).

For the comparison of immunofluorescence signals of α-catenin and α18 antibody staining, a median filter (2 × 2 pixels) was used to reduce the image noise, and immunofluorescence signals in each channel were normalized to total signal strength in each image. The signal intensity ratio between α-catenin and α18 antibody staining was standardized based on the average ratio in each image.

**Biotinylation of apical surface proteins in the mouse uterus and protein extraction**

Uteri were dissected from non-pregnant or E4.5 mice, washed in PBS, and placed on Kimwipes moistened with PBS. Then, 100 μL of 2.5 mM sulfo-NHS-biotin (A39256, Thermo Fisher Scientific) dissolved in PBS was injected into the uterus from near the uterine tubal junction using a 30-gauge needle. The injected uteri were incubated for 20 min at room temperature and then washed in TBS to quench. The uteri were cut into several pieces and smashed in a disposable homogenizer in PBS. After centrifugation at 20,000 × g for 10 min, the supernatant was kept as S0, and the pellet was resuspended in RIPA buffer followed by centrifugation. The supernatant was kept as S1. Both S0 and S1 protein extracts were analyzed as described below.

**Purification of biotinylated peptides for mass spectrometry**

The extracted proteins were precipitated using the methanol/chloroform method (*30*) (*31*), and then dissolved in 20 mM triethylammonium bicarbonate (TEAB) buffer pH 8.5. The extracts were incubated with Tris (2-carboxyethyl) phosphine (TCEP) to a final concentration of 20 mM at 55°C for 30 min, followed by incubation with chloroacetamide to a final concentration of 40 mM at RT for 30 min in the dark. Then, 10 µg of trypsin/LysC mix, MS grade (V5071, Promega) was added to 1 mg of protein, and the samples were incubated overnight at 37°C. The samples were mixed with 20 µL of Tamavidin-2 (136-18341, Fuji Film WAKO) and tumbled at 4ºC for 2 h. The beads were washed with PBS, and peptides bound to the beads were eluted by incubating the beads with 2 mM biotin for 15 min at 37°C. The eluates were purified using C-18 spin columns (89870, Thermo Fisher Scientific) and used for mass spectrometry analysis.

**Mass spectrometry analysis**

Nano-LC–MS analysis was performed using an EASY-nLC 1000 (Thermo Fisher Scientific) connected to an Orbitrap Elite mass spectrometer (Thermo Fisher Scientific). Small protein–containing and peptide fragment–containing fractions were loaded onto a trapping column (75 µm i.d. × 20 mm, Acclaim PepMap 100 C18 LC; Thermo Fisher Scientific). Peptides were subsequently eluted from the trapping column and separated on a nanocapillary column (75 µm i.d. × 125 mm, NTCC-360; Nikkyo Technos) with a 30-min 5–35% acetonitrile (containing 0.1% formic acid) gradient, followed by a 2-min 35–80% gradient, with a final 8-min isocratic step at 80% acetonitrile and a flow rate of 300 nL/min. The nano-HPLC eluate was introduced into a mass spectrometer via an electrospray ionization (ESI) interface at a spray voltage of 2.0 kV. The mass spectrometer was operated in positive ion mode with a capillary temperature of 250ºC. Mass spectra were obtained by scanning from m/z 350 to m/z 2000. Nano-LC–MS/MS analysis was performed by selecting the indicated molecular ion as the precursor ion at 30% normalized collision energy using the higher energy collision dissociation mode. The acquired MS/MS spectra were analyzed with MASCOT version 2.8.0.1 using the mouse protein database. The database search was performed using a precursor ion mass tolerance of 10 ppm and a fragment ion mass tolerance of 0.1 Da. For the identification of small proteins, database searches were limited to fully tryptic peptides with two missed cleavages allowed; carbamidomethyl cysteine, biotinylated lysine, and biotinylated N-terminus were set as variable modifications (FDR $\leq$ 0.01). Datasets of MS/MS spectra were deposited in the Japan Proteome Standards Repository/Database (jPOST; https://repository.jpostdb.org) (*32*). The accession numbers are PXD044534 for ProteomeXchange and JPST002275 for jPOST

**C-CPE production in bacteria and injection into the mouse uterus**

A cDNA encoding C-CPE (COOH-terminal half fragment of *Clostridium perfringens* enterotoxin, 184–319 amino acids) fused to GST (GST-C-CPE) was used (*33*) . The GST-C-CPE cDNA was ligated into the bacterial expression vector pCold II (3362, Takara Bio Inc,), and a 6-His tag was added to the amino terminus of GST and designated pCold-C-CPE. For a negative control, GST with 6-His tag was ligated into pCold II and designated pCold-GST. The plasmids were introduced into BL21(DE3)-pLysS competent cells (L1195, Promega). The transformants were cultured in 0.5 mL of LB medium containing 50 μg/mL ampicillin and 34 μg/mL chloramphenicol overnight at 37ºC. The cultured bacteria were inoculated into 150 mL of the same medium, shaken for 1 h, and then cooled to 18ºC for 1 h. After adding IPTG (isopropyl β-D-thiogalactopyranoside, final concentration, 1 mM), the bacteria were incubated at 18°C overnight.

The bacteria were collected by centrifugation and lysed by freezing, thawing, and sonication in 1 mL of PBS containing 0.1% Triton-X100 and a protease inhibitor mix. The lysate was centrifuged for 10 min at 20,380 x g. The supernatant was mixed with 100 μL of Talon Metal Affinity Resin (635501, Takara Bio Inc) and rotated for 2 h at 4°C. Then, the resin was washed four times with PBS, and proteins bound to the resin were eluted with PBS containing 0.5 M imidazole. The eluate was dialyzed in PBS overnight.

Mice at the E3.75 were anesthetized with a triple anesthetic [medetomidine (0.75 mg), midazolam (4 mg), and butorphanol (5 mg), 0.1 mL/10 g weight]. Approximately 1 cm long cuts were made at the left side of the back skin and the inside muscle layer with scissors. One of the terminal regions of the uterus close to the oviduct was pierced with a 27-gauge needle, and 10 μL of C-CPE or GST protein in PBS (1 μg/μL) was injected into the uterine lumen with a microcapillary. The wound site was closed with a surgical stapler. The uterus was dissected out at E4.5, embedded in OCT compound, frozen in liquid nitrogen, and sectioned at 16 μm thickness. The sections were fixed in cold methanol for 20 min, dried, and immunostained as described above. The tissue sections for immunohistochemistry were selected randomly and analyzed the signal localization or effect of C-CPE treatment. Statistics, two-sided Student's t-test in 95% confidence interval. Centre line; median, box edge; upper and lower quartiles, whiskers; 1.5 x interquartile range. N = 2 mice for GST and 3 mice for C-CPE. These analyses were conducted with R.

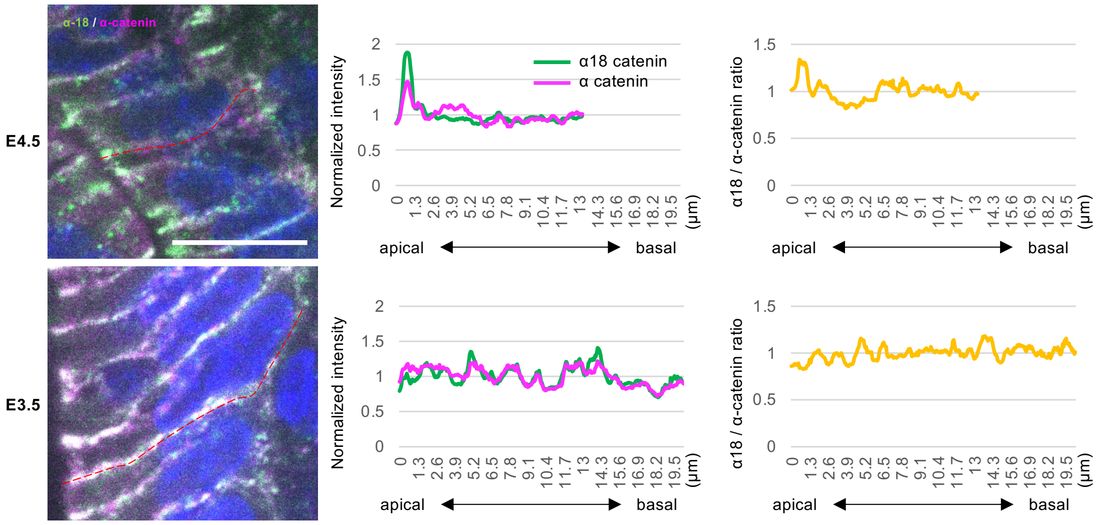


**Fig. S1. Comparison of α-catenin and α18 signal intensities between the open LE at E3.5 and the closed LE at E4.5.**

Immunofluorescence detection of α-catenin and α18. The intensity profiles of representative lateral cell borders (red dot lines) along the apical–basal axis are shown in the middle panels. The normalized ratio between α-catenin and α18 signals is shown in the right panels. Scale bars: 10 μm

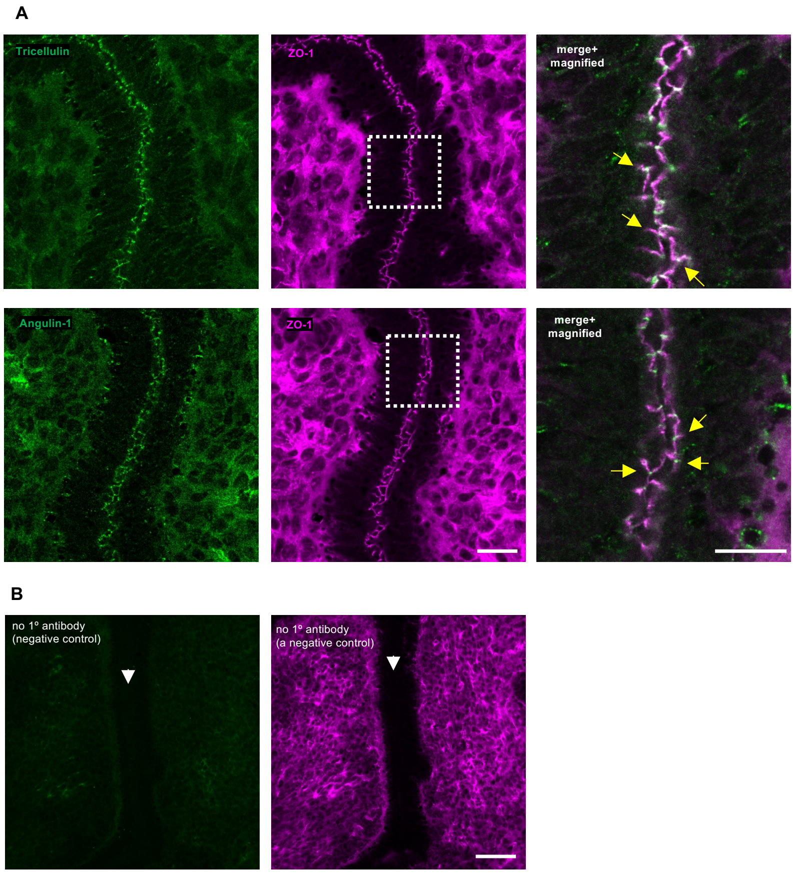


**Fig. S2. Immunofluorescence detection of tricellulin and angulin-1.**

(**A**) Immunostaining of tricellulin and angulin-1 co-stained with ZO-1 in a transverse uterine section at E4.5. In the merged and magnified images, the triple bifurcation of the junction is indicated by yellow arrows. Scale bars: 50 µm in the left and middle panels and 20 µm in the right magnified panels. (**B**) Signals detected in the negative control samples. Secondary antibody staining detected cytoplasmic signals as the background. The apical contact site is indicated with white arrows.

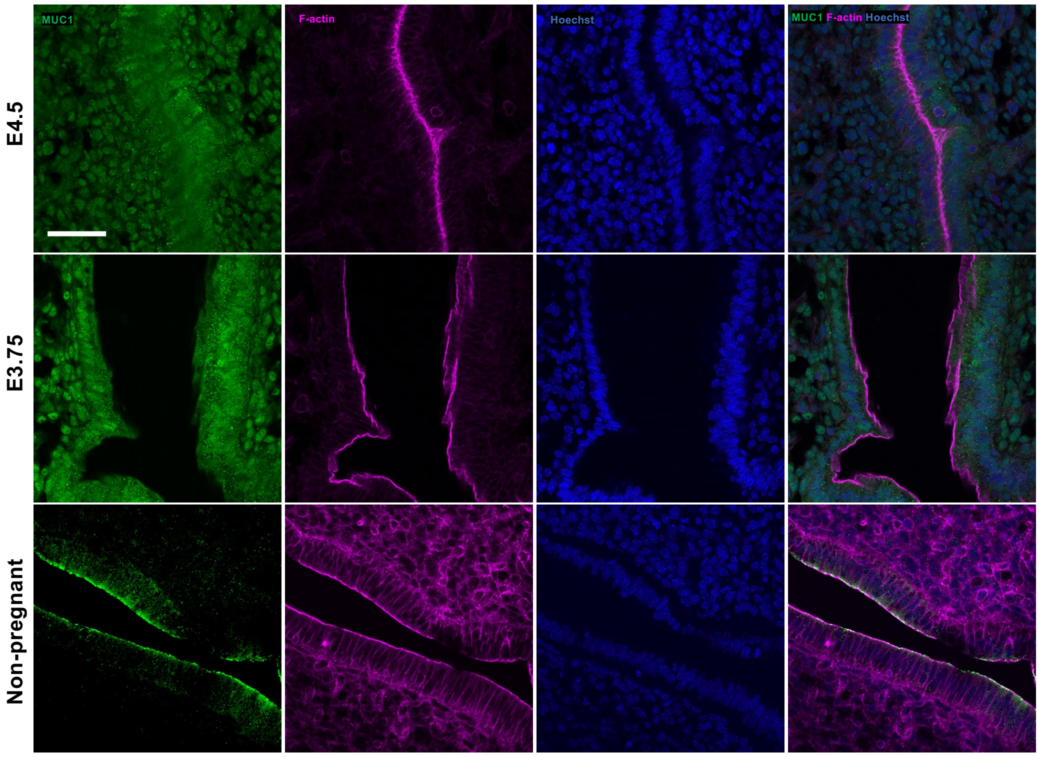


**Fig. S3. Mucin-1 decreases in the apical region of the LE after pregnancy.**

Immunostaining of mucin-1 in the transverse uterine sections at E4.5, E3.75, and non-pregnant samples. F-actin and nuclei are visualized with phalloidin and Hoechst 33342, respectively. Scale bars: 50 µm

**Movie S1.** Expansion of uterine tissue after application of an external force with forceps as seen in both sides of the image.
